# Induced Alpha And Beta Electroencephalographic Rhythms Covary With Single-Trial Speech Intelligibility In Competition

**DOI:** 10.1101/2022.12.31.522365

**Authors:** Vibha Viswanathan, Hari M. Bharadwaj, Michael G. Heinz, Barbara G. Shinn-Cunningham

**Affiliations:** Neuroscience Institute, Carnegie Mellon University, Pitttsburgh, PA 15213; Department of Communication Science and Disorders, University of Pittsburgh, Pitttsburgh, PA 15260; Department of Speech, Language, and Hearing Sciences, Purdue University, West Lafayette, IN 47907

## Abstract

Neurophysiological studies suggest that intrinsic brain oscillations influence sensory processing, especially of rhythmic stimuli like speech. Prior work suggests that brain rhythms may mediate perceptual grouping and selective attention to speech amidst competing sound, as well as more linguistic aspects of speech processing like predictive coding. However, we know of no prior studies that have directly tested, at the single-trial level, whether brain oscillations relate to speech-in-noise outcomes. Here, we combined electroencephalography while simultaneously measuring intelligibility of spoken sentences amidst two different interfering sounds: multi-talker babble or speech-shaped noise. We find that induced parieto-occipital alpha (7–15 Hz; thought to modulate attentional focus) and frontal beta (13–30 Hz; associated with maintenance of the current sensorimotor state and predictive coding) oscillations covary with trial-wise percent-correct scores; importantly, alpha and beta power provide significant independent contributions to predicting single-trial behavioral outcomes. These results can inform models of speech processing and guide noninvasive measures to index different neural processes that together support complex listening.

## 2. Introduction

Understanding speech in the presence of interfering sounds—e.g., competing talkers or other sources of background noise—is a difficult perceptual task that our brains solve everyday [1]. However, the neural mechanisms facilitating “cocktail-party” listening remain poorly understood. Neurophysiological studies in humans [using electro-(EEG) and magneto-encephalography (MEG) as well as invasive intracranial recordings] and other animal species suggest that brain rhythms [2] in different frequency bands may mediate sensory processing [3]. This, in turn, may facilitate speech understanding in competition. For instance, in a mixture of competing sources, brain oscillations in the low-frequency delta (1–3 Hz) and theta (3–7 Hz) bands preferentially phase-lock to the slow temporal fluctuations (i.e., envelopes) in attended speech [4, 5], while power fluctuations in the low-gamma (30–70 Hz) [6] and high-gamma (70–120 Hz) [7, 8] bands selectively synchronize to attended versus ignored speech envelopes. This target-speech envelope phase-locking in the brain may aid listeners in selectively processing a target speech source in an acoustic mixture, thereby influencing speech intelligibility across different everyday listening conditions [9].

In addition to phase-locked (or evoked) neuronal oscillations, induced brain rhythms have also been implicated in cocktail-party listening. For instance, focused auditory attention leads to an increase in the power of the alpha (7–15 Hz) rhythm in parieto-occipital areas, which is specifically thought to be a hallmark of neuronal mechanisms related to suppression of sensory distractors [10–14] like visual input [15]. During auditory spatial selective attention, parieto-occipital alpha power becomes lateralized: alpha power increases contralateral to the hemifield of distracting sounds (i.e., ipsilateral to the hemifield of focus) [16–18]. This alpha lateralization has been reported to predict individual differences in spoken-digit identification when listeners hear a mixture of spatially separated sources [19]. Moreover, even for tasks that involve spatially co-localized speech and distractor sources, prior studies report positive correlation between the overall magnitude (versus lateralization) of alpha power in centro-parietal EEG channels and speech comprehension across signal-to-noise ratios (SNRs; for spoken sentences) [20] and across individuals (for spoken digits) [21].

In contrast to induced alpha, which has been implicated in auditory attention, prior work suggests that the beta (13–30 Hz) rhythm may relate to maintenance of the current sensorimotor state [22] and sensorimotor predictive coding [23, 24]. More generally, motor-theory accounts of speech recognition posit that sensorimotor integration between fronto-motor areas controlling articulation (e.g., inferior frontal gyrus and premotor cortex) and temporal-parietal cortical areas implicated in phonetic category representation mediates top-down sensory prediction to modulate and stabilize speech representation [25–27], especially in adverse listening conditions such as in background noise [28, 29]. In line with this notion, frontal beta power correlates with sensory prediction precision in vocoded word identification [30], with auditory cortical entrainment to continuous speech [31, 32], and with comprehension for time-compressed speech sentences [33]. Moreover, across individuals, beta-band synchrony between premotor and temporal-parietal cortical regions correlates positively with syllable identification in noise [34]. Finally, some studies found beta power to be greater at the target word for syntactically and semantically legal sentences compared to sentences containing a syntactic or semantic violation [35–38].

Despite the prior literature linking alpha and beta rhythms to speech processing, we know of no prior studies that tested whether trial-to-trial variations in the overall magnitude of induced parieto-occipital alpha power and frontal beta power relate to trial-wise speech intelligibility when competing sounds are present. The present study explored this question using human EEG and simultaneous speech intelligibility measurements of spoken sentences under masking.

Because different competing sounds could drive different degrees of demand on selective attention versus contextual prediction (e.g., attentional demand—and hence alpha power—may be greater when the masker is a competing speech stream or multi-talker babble versus stationary noise; [39]), we used two different maskers in this study: multi-talker babble and speech-shaped stationary noise. We examined the extent to which the overall magnitude of induced oscillatory power in different frequency bands relates to speech intelligibility in each masking condition on a trial-by-trial basis. Building on our previous work, where we related phase-locked neural responses to speech understanding in different listening conditions in the same dataset [9], here we focused on induced brain activity. Specifically, we examined frequency bands in which the prior literature reports induced responses to speech. Because we sought to quantify induced activity on a single-trial level, we focused on alpha and beta rhythms here (rather than higher-frequency gamma band activity) due to the relatively greater SNR of alpha and beta in EEG measurements [40, 6].

## 3. Materials and Methods

The stimuli, participants, experimental design, and hardware used in the current study are described in detail in the materials and methods in our prior work [9]. Below, we describe each briefly.

### 3.1. Stimulus generation

Target speech that listeners were instructed to attend were Harvard/Institute of Electrical and Electronics Engineers (IEEE) sentences [41] spoken in a female voice and recorded as part of the PN/NC corpus [42]. Stimuli were created for two different speech-in-noise experimental conditions, as described below.

1. Condition 1: Speech in babble (SiB). Speech was added to spectrally matched four-talker babble at -2 dB SNR. The long-term spectra of the target speech sentences were adjusted to match the average (across instances) long-term spectrum of four-talker babble. In creating each SiB stimulus, a babble sample was randomly selected from a list comprising 72 different four-talker babble maskers obtained from the QuickSIN corpus [43]. All four talkers in each four-talker babble masker used are adults; each babble sample had a distribution of 3 female talkers and 1 male talker.
2. Condition 2: Speech in speech-shaped stationary noise (SiSSN). Speech was added to spectrally matched stationary Gaussian noise, i.e., speech-shaped stationary noise, at -5 dB SNR. The long-term spectra of the target speech sentences and that of stationary noise were adjusted to match the average (across instances) long-term spectrum of four-talker babble. A different realization of stationary noise was used for each SiSSN stimulus.

The particular SNRs used for SiB and SiSSN yielded average speech intelligibility values close to 50% (Figure 1; also see [9]), which helped avoid floor and ceiling effects when quantifying percent-correct scores on a trial-by-trial basis. Note that the speech and masker sources in each acoustic mixture were co-localized (presented diotically) for all stimuli.

**Figure 1.**
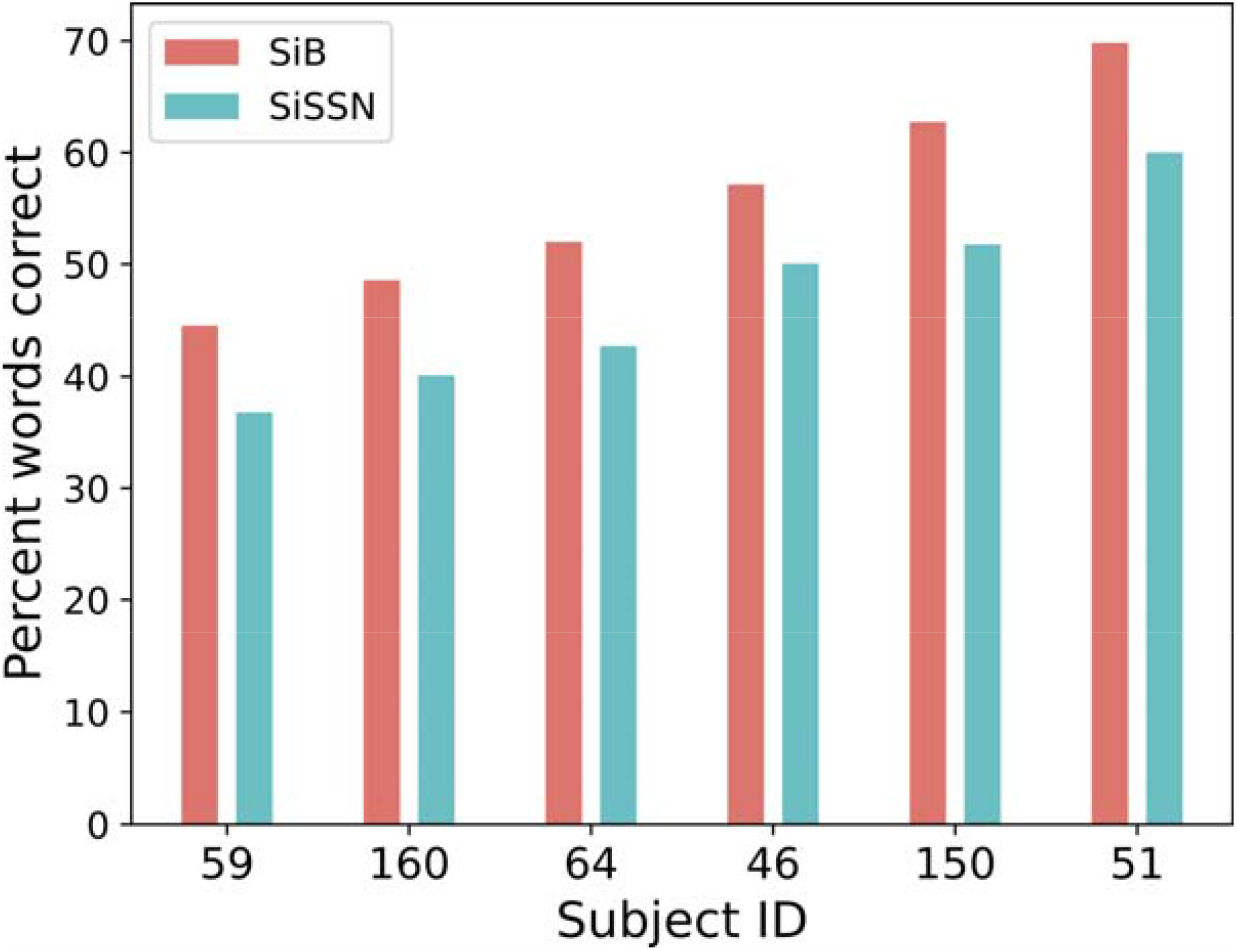
Percent keywords correct as a function of subject and experimental condition (SiB versus SiSSN). Data shown are pooled over trials, and sorted along the x-axis according to average behavioral performance across the two conditions.

### 3.2. Participants

Data from six human subjects (one male, five female) aged 19–31 years were analyzed for this study. All subjects were native speakers of North American English, had pure-tone hearing thresholds better than 20 dB hearing level in both ears at standard audiometric frequencies between 250 Hz and 8 kHz, and reported no history of neurological disorders. All subjects also had distortion-product and click-evoked otoacoustic emissions within the normal range [44] as well as normal tympanograms. All human subject measures were conducted in accordance with protocols approved by the Purdue University Institutional Review Board and the Human Research Protection Program, and all subjects provided informed consent. Data were collected from each subject over the course of one or two visits (with a total visit time of approximately 5 hours).

### 3.3. Experimental design

Each subject performed 175 trials of speech intelligibility testing for each of the two experimental conditions, with a distinct target sentence in every trial. In total, 1050 trials were collected for each experimental condition across the subject cohort. The different experimental conditions were intermingled across trials.

Thirty-two-channel EEG was measured as subjects performed the speech identification task. The target speech sentences were presented at a sound level of 72 dB sound pressure level (SPL), while the level of the background was set to obtain the desired SNR.

Subjects were instructed that they would be listening for a woman’s voice speaking a sentence in each trial and that at the end of the trial they would have to verbally repeat the sentence back to the experimenter sitting beside them. They were told that it would be the same woman’s voice every time but that the type and level of background noise would vary across trials. They were also told that the noise would start first in each trial with the target woman’s voice starting approximately one second later. They were encouraged to guess as many words as they could if they heard a sentence only partially.

At the beginning of each trial, subjects were presented with a visual cue that read “stay still and listen now” in red font. The audio stimulus started playing one second afterward. In every trial, the background noise started first, while the target speech started 1.25 seconds later to allow sufficient time to cue the subjects’ attention to the stimulus. The target was at least 2.5 seconds long. After the target sentence ended, the background noise continued for a short, variable amount of time. Two hundred ms after the noise ended, subjects were presented with a different visual cue that read “repeat now” in green font, cueing them to report the target sentence. This delayed response design avoided motor artifacts and speech-motor preparation signals from contributing to the EEG recorded during listening. For each trial, intelligibility was scored on five pre-determined keywords (which excluded articles and prepositions) in the target sentence and then converted to a percent-correct score (whose value in each trial was either 0, 20, 40, 60, 80, or 100). Mistakes in plurality (e.g., “cats” versus “cat”) and tense (e.g., “relax” versus “relaxed”) were not penalized. The role of the experimenter sitting in the sound booth (“sitting experimenter”) was to check off the specific keywords (out of the five in each sentence) that were answered correctly; full responses from subjects were not noted. Two different sitting experimenters were used in the study. Before the actual EEG experiment, subjects performed a short training demo task, which used the same listening conditions and target speaker’s voice as the actual experiment but a different set of Harvard/IEEE target sentences from the main experiment. Note that our experimental paradigm is in line with the guidelines proposed in Wöstmann et al. [45].

### 3.4. Hardware

The entire experiment was conducted in a sound-treated booth. A personal desktop computer controlled all aspects of the experiment, including triggering sound delivery and storing data. Special-purpose sound-control hardware (System 3 real-time signal processing system, including digital-to-analog conversion and amplification; Tucker Davis Technologies, Alachua, FL) presented audio through insert earphones (ER-2; Etymotic, Elk Grove Village, IL) coupled to foam ear tips. The earphones were custom shielded by wrapping the transducers in layers of magnetic shielding tape made from an amorphous cobalt alloy (MCF5; YSHIELD GmbH & Co., Ruhstorf, Germany) and then placing them in 3-mm-thick aluminum enclosures to attenuate electromagnetic interference. The signal cables driving the transducers were shielded with braided metallic Techflex (Techflex, Sparta, NJ). All shielding layers were grounded to the chassis of the digital-to-analog (D/A) converter. The absence of measurable electromagnetic artifact was verified by running intense click stimuli through the transducers with the transducers positioned in the same location relative to the EEG cap as actual measurements but with foam tips left outside the ear. All audio signals were digitized at a sampling rate of 48.828 kHz. The EEG signals were recorded at a sampling rate of 4.096 kHz using a BioSemi (Amsterdam, The Netherlands) ActiveTwo system. Recordings were done with 32 cephalic electrodes and two additional earlobe electrodes.

### 3.5. EEG processing

All six subjects were able to stay still during the presentation of the sentences and respond on cue. EEG signals were preprocessed by re-referencing channel data to the average of the two earlobe reference electrodes. Then, the signal space projection method was used to construct spatial filters to remove eye blink and saccade artifacts [46]. Finally, the broadband EEG was bandpass filtered between 1 and 400 Hz and parceled into epochs, each of which corresponded to a distinct trial. Each epoch started 0.5 seconds before the “stay still and listen now” cue of the corresponding trial, and ended 2.5 seconds after the onset of the target sentence; thus the total duration of each epoch was 5.25 seconds. This epoch period included all of the five keywords on which participants were scored for every sentence.

For each subject and experimental condition, the EEG response spectrogram in each epoch was calculated using a Slepian-tapered complex exponential wavelet (which minimizes spectral leakage, i.e., any sidelobe energy that may bias spectrogram estimates) [47, 48]. Five cycles were used to estimate each time-frequency bin with a time-full-bandwidth product of 2. Note that the average EEG time course across epochs was not subtracted prior to computing the single-trial spectrograms. Because the stimuli contained different naturally uttered speech sentences across different trials, the trial-averaged EEG time course is negligibly small compared to the induced power captured by the spectrograms.

Based on our observation of clear induced brain oscillations in the alpha (7–15 Hz) and beta (13–30 Hz) bands (Figure 2; the frequency ranges considered for alpha and beta are based on prior literature [6, 10, 13, 17–19, 23, 24, 31]), we derived scalp topomaps in each of these two bands for the pre-stimulus (the one-second-long time period between the “stay still and listen now” cue and the stimulus presentation) and during-stimulus (the 3.75-seconds-long time period between the start of stimulus presentation and end of the target sentence) periods; this was done by averaging the response spectrogram over all band-specific frequencies, epochs, subjects, experimental conditions, and time samples in the corresponding period.

**Figure 2.**
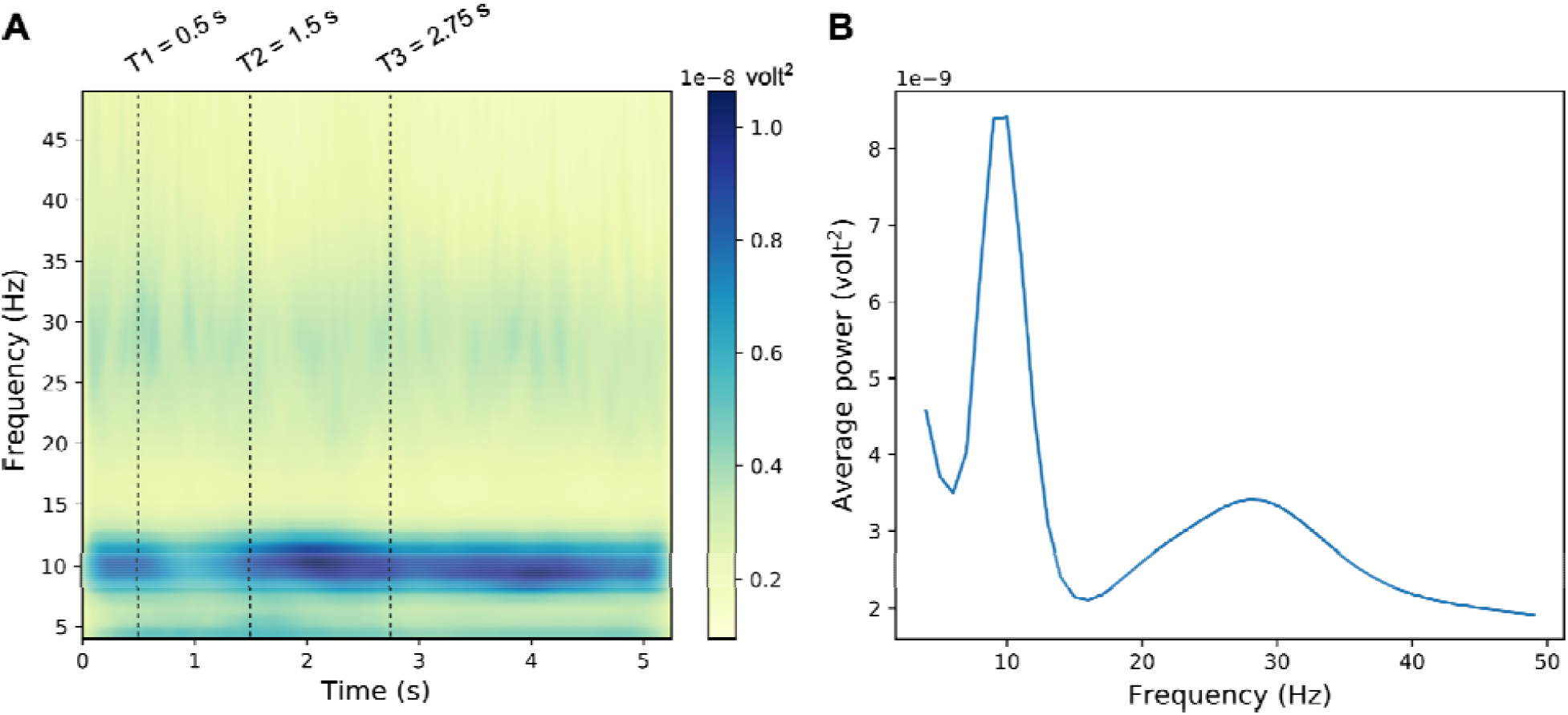
Average EEG response spectrogram (A) and spectrum (B). The spectrogram and spectrum shown are averaged over the 32 EEG channels, and all trials, subjects, and experimental conditions. Note that the time before T1 corresponds to baseline. At T1, the “stay still and listen now” visual cue was shown. At T2, the audio stimulus started playing. At T3, presentation of the target speech sentence started; target presentation lasted until at least 5.25 s (and was longer for the longer sentences).

To obtain overall measures of alpha power in each trial, the spectrogram in the corresponding epoch was averaged in the alpha band over parieto-occipital channels (A9, A10, A11, A12, A13, A14, A15, A16, A17, A18, A19, A20, A21, and A22; based on Figure 3A topomap). This was done separately over the pre-stimulus period and the during-stimulus period. An overall measure of trial-specific pre- and during-stimulus beta-band power was obtained using a similar approach, but by using frontal channels (A1, A2, A3, A4, A5, A6, A25, A26, A27, A28, A29, A30, and A31) instead (based on Figure 3B topomap). Since alpha power in the pre- and during-stimulus periods were strongly correlated (Figure 4A), we combined the pre- and during-stimulus power by averaging before performing further analysis. Similarly, since the beta power in the pre- and during-stimulus periods were correlated (Figure 4B), the average power across pre- and during-stimulus periods was used for all further analysis.

**Figure 3.**
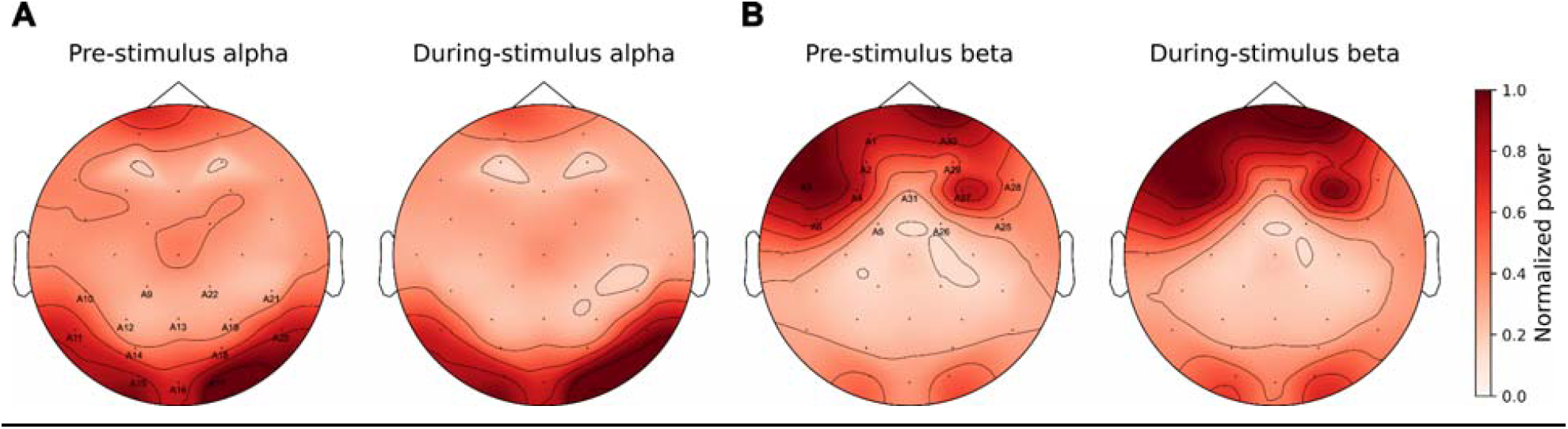
Average scalp topography maps for the induced oscillatory power in the alpha (A) and beta (B) bands, shown separately for the pre- and during-stimulus time periods. The topomaps are averaged over band-specific frequencies, trials, subjects, experimental conditions, and time samples. Parieto-occipital channels (A9, A10, A11, A12, A13, A14, A15, A16, A17, A18, A19, A20, A21, and A22; where alpha power is strongest) and frontal channels (A1, A2, A3, A4, A5, A6, A25, A26, A27, A28, A29, A30, and A31; where beta power is strongest) are indicated in Panels A and B, respectively.

**Figure 4.**
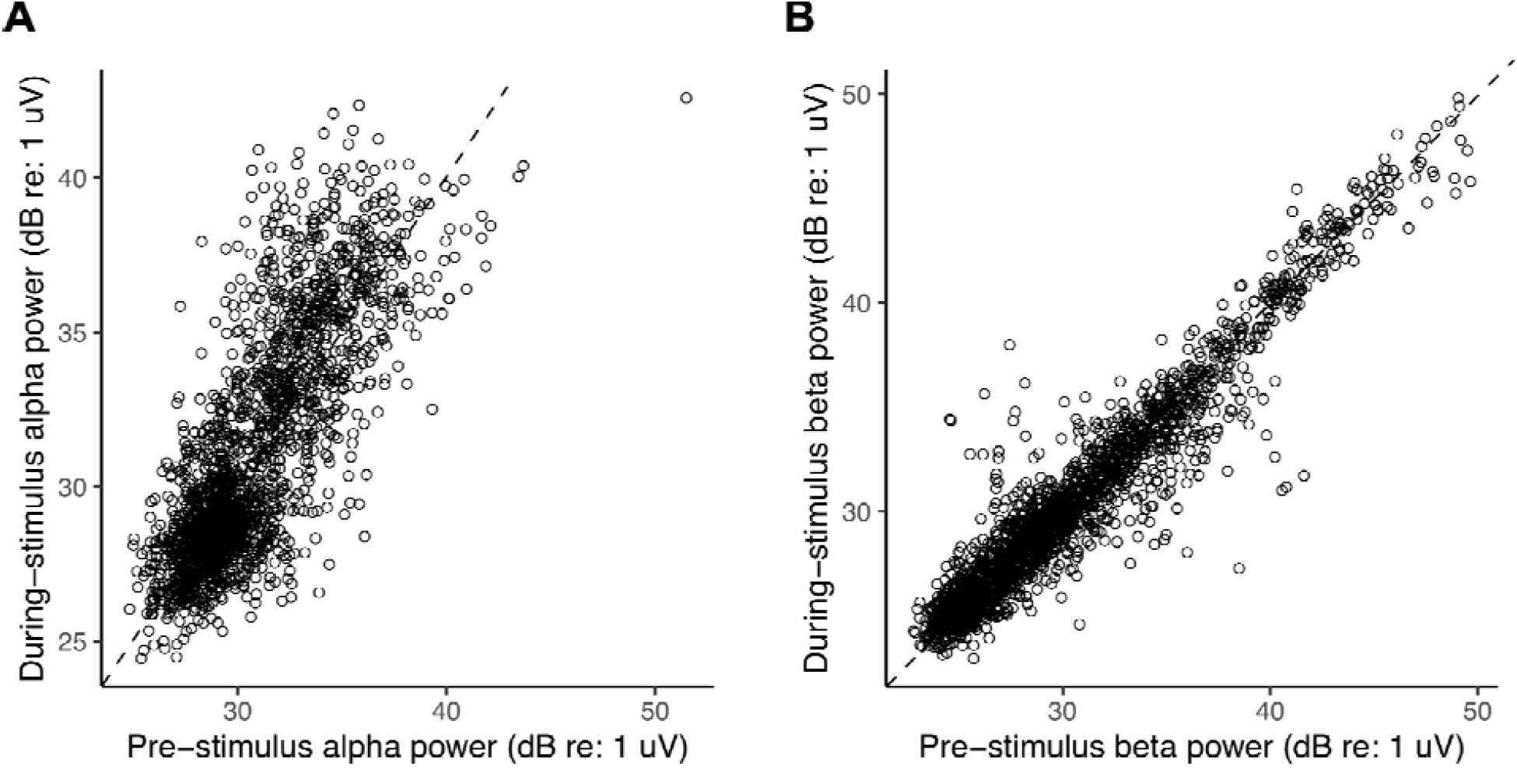
Overall parieto-occipital alpha (A) and frontal beta (B) power in different trials (across subjects and conditions) in the pre-stimulus period against the corresponding values in the during-stimulus period. The dashed line in each plot indicates points where pre- and during-stimulus power are equal. Note that the unit for alpha and beta power is dB relative to one microvolt, calculated as 10log_10_(power/10^-12^).

### 3.6. Statistical analysis

We tested whether greater alpha power is associated with higher percent-correct score across different trials. A linear model was built with the different trials across subjects as observations. The log of alpha power was taken as the response variable (to ensure response data are normally distributed; [49]) while percent-correct score and condition (SiB versus SiSSN) were predictors. Percent correct was treated as an ordered factor variable (with six levels: 0, 20, 40, 60, 80, and 100) and condition as a factor variable (with two levels). Statistical tests used ANOVA (Type II tests with the F statistic) along with post-hoc t-tests. The same approach was used to test whether greater beta power is associated with higher percent-correct score across trials.

We performed several posthoc analyses to quantitatively assess the precise relationship between percent-correct score and alpha or beta power. First, we examined the individual terms in the omnibus model to assess the respective contributions of the linear, quadratic, cubic, and fourth order terms in the model. The linear term had the largest contribution to the overall main effect in both the alpha and beta models. To understand the linear term further, we performed pairwise t-tests for percent-correct scores going from 0 to 20, 20 to 40, 40 to 60, etc. (i.e., successive difference contrast coding).

To test whether alpha and beta power are correlated across trials, we used a linear mixed-effects model to account for any random effect of subject. Log beta power in different trials across subjects was the response, and log alpha power, condition (factor variable with two levels), and subject were predictors. Anova (Type II Wald F tests with Kenward-Roger degree of freedom; [50]) was used for statistical testing.

To test whether there are individual differences in the overall magnitude of alpha power, overall magnitude of beta power, or the alpha-to-beta power ratio across trials, we constructed linear models with either log alpha power, log beta power, or their ratio as the outcome, and with subject (factor variable with six levels) and condition (factor variable with two levels) as fixed-effect predictors. ANOVA (Type II tests with the F statistic) was used for statistical testing.

Behavioral outcomes across different trials may be influenced by top-down effects like selective attention and contextual prediction. To explore this possibility, we examined response error patterns, computing the histogram (across trials and subjects) of number of keywords correct per sentence separately for each experimental condition. If either selective attention or predictive coding—or a combination of the two—influence trial-wise behavioral outcome, performance across the different keywords should be correlated within a trial. We tested for this effect separately for each experimental condition by comparing the histogram of number of keywords correct per sentence against the distribution under the null hypothesis of independent outcomes across different keywords. Under the null hypothesis, the performance on any particular keyword in a sentence (i.e., whether or not the word was reported correctly) has a Bernouilli distribution with parameter p = average proportion correct score for the particular condition. Moreover, the probability of reporting correctly *x* keywords out of a total of 5 keywords per sentence is binomial with parameters n = 5, and p = average proportion correct score for the condition. Assuming independent outcomes across different sentences, the probability that *M* sentences out of a total of 1050 sentences per condition (pooled over all six subjects; each subject performed 175 sentences per condition) had *x* keywords correct is also binomial, with parameters n = 1050 and p = probability that *x* keywords per sentence are correct. We compared this final probability distribution (which models the null distribution of independent keyword outcomes in each condition) with the histogram of number of keywords correct per sentence. Specifically, we generated p-values describing, for each experimental condition, the likelihood of observing the actual correlation across keywords within each trial, assuming that performance on the words was truly independent.

We used a multinomial linear regression model to test whether beta power contributes additionally to predicting percent-correct score over the contribution of alpha power alone, and vice-versa (i.e., whether alpha power contributes additional predictive power over that contributed by beta power alone). The percent-correct score in different trials across subjects was the response (treated as an ordered factor variable with six levels: 0, 20, 40, 60, 80, and 100); the predictors were log alpha power, log beta power, and condition (factor variable with two levels). Likelihood-ratio Type II tests were used for statistical testing by calculating the deviance (i.e., -2 times log likelihood-ratio) and comparing it to a chi-squared distribution [51].

### 3.7. Software accessibility

Stimulus presentation was controlled using custom MATLAB (The MathWorks, Inc., Natick, MA) routines. EEG preprocessing was performed using the open-source software tools MNE-PYTHON [52] and SNAPsoftware [53]. All further data analyses were performed using custom software in PYTHON (Python Software Foundation, Wilmington, DE). Statistical analyses were performed using R (R Core Team; www.R-project.org). Visualizations used the colorblind-friendly Colorbrewer [54] and Color Universal Design [55] colormap palettes. Our custom code is publicly available on https://github.com/vibhaviswana/inducedOscillationsAndSpeechIntelligibility.

### 3.8. Data availability

The datasets used in the current study are available from the corresponding author on reasonable request.

## 4. Results

We wished to quantify induced brain oscillations in different canonical frequency bands [2] on a trial-by-trial basis and relate those to speech intelligibility, also measured on a trial-by-trial basis. For this, we first computed the EEG response spectrogram in each trial (see Materials and Methods: EEG processing) to examine induced (versus evoked) oscillatory activity in different frequency bands. Figure 2A shows the average EEG spectrogram over the 32 EEG channels, and all trials, subjects, and experimental conditions. Figure 2B shows the average EEG spectrum obtained by computing the mean over time of the spectrogram shown in Figure 2A. As seen in Figure 2, induced brain oscillations are clearly visible in the alpha (7–15 Hz) and beta (13–30 Hz) bands. Because we did not see any induced activity outside the alpha and beta frequency ranges, we restricted all further analyses in the current study to just the alpha and beta bands.

To better understand the neural sources of these alpha and beta induced oscillations, we computed their scalp topography (see Materials and Methods: EEG processing). Figures 3A and 3B show average (over band-specific frequencies, trials, subjects, experimental conditions, and time samples) scalp topomaps for the alpha and beta bands, respectively; the topomaps are plotted separately for the pre- and during-stimulus periods so as to be able to visualize any differences in the contributions of preparatory rhythmic activity and stimulus-induced oscillatory activity [10]. Results suggest that the strongest alpha power occurs in the parieto-occipital channels (Figure 3A) and the strongest beta power occurs in the frontal channels (Figure 3B).

Based on Figure 3 scalp topomaps, we used parieto-occipital EEG channels to derive an overall measure of alpha power for each trial from the EEG response spectrogram; we did this separately for the pre- and during-stimulus periods (see Materials and Methods: EEG processing). Similarly, we derived an overall measure of trial-specific pre- and during-stimulus beta power, but by using frontal EEG channels instead. Figures 4A and 4B plot the power in different trials across subjects and conditions in the pre-stimulus period against the corresponding values in the during-stimulus period for the alpha and beta bands, respectively. Across trials, pre- and during-stimulus induced oscillation power was significantly correlated for both alpha (R^2^ = 0.6292, DF = 2098, p < 2e-16) and beta (R^2^ = 0.9151, DF = 2098, p < 2e-16). For beta, the during-stimulus power was roughly equal to the pre-stimulus power (in Figure 4B, data fall along the identity line); however, for alpha, the during-stimulus power was consistently higher than the pre-stimulus power (in Figure 4A, data fall above the diagonal), but the pre-stimulus value nonetheless predicted the during-stimulus alpha power value. We were interested in whether the difference in alpha (and beta) power across trials was related to behavioral performance and wished to quantify the level of alpha (and beta) in each trial relative to the other trials. Therefore, we averaged the pre- and during-stimulus periods together for all further analyses.

Across trials, we compared percent-correct score with the corresponding alpha and beta power. Figure 5A shows alpha power versus percent-correct score in different trials across subjects, separately for each condition. Alpha power covaried significantly with percent-correct score within condition [F(5,2093) = 4.7789, p = 0.0002397; see Materials and Methods: Statistical analysis for details]. Figure 5B shows beta power versus percent-correct score in different trials across subjects, separately for each condition. Beta power too covaried significantly with percent-correct score within condition [F(5,2093) = 6.4915, p = 5.346e-06]. Posthoc analyses revealed that the largest contribution to the main effect of score on alpha and beta power came from the linear term, indicating that alpha and beta power increased with score (T = 4.216, p = 2.59e-05 for alpha; T = 4.173, p = 3.13e-05 for beta). The quadratic term was also significant in predicting alpha power, but carried a negative coefficient (T = -2.521, p = 0.0118) in line with the plateauing of alpha power with increasing score seen in Figure 5A. In the model for beta power, the cubic term was also significant (T = 3.003, p = 0.00271), in line with the U-shaped trend seen over the 20–100% range of scores in Figure 5B. We conducted pairwise t-tests to compare the changes in alpha and beta power for each step increase in percent-correct score. These sequential-difference-contrast analyses showed that alpha and beta power increased when percent-correct score increased from 0 to 20 (T = 1.994, p = 0.0463 for alpha; T = 4.249, p = 2.24e-05 for beta); however, for the successive steps (20 to 40, 40 to 60, etc.), the increase in power was not significant for either alpha or beta. Figure 5A also suggests that for a given percent-correct score, alpha power is greater for speech in stationary noise than in babble; however, statistical testing did not reveal a significant effect of experimental condition (SiB versus SiSSN) on alpha power. Moreover, when we included the interaction between percent-correct score and condition (in addition to the main effects of these two variables) as a predictor in the linear model with alpha or beta power as the outcome, statistical testing did not reveal a significant effect of the interaction between percent-correct score and condition on either alpha or beta power.

**Figure 5.**
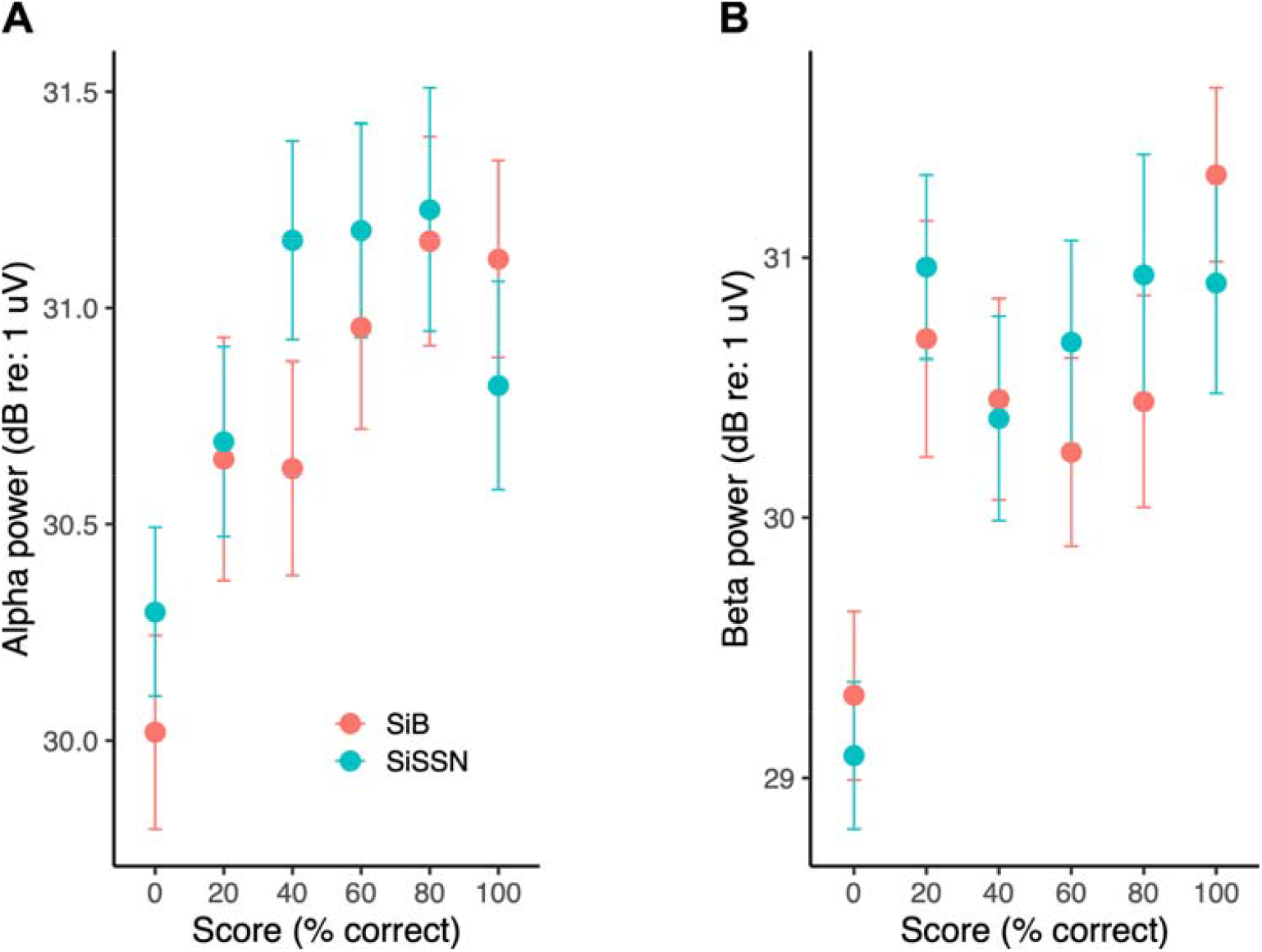
Alpha (A) and beta (B) power [mean and standard error of the mean (STE)] versus percent-correct score in different trials across subjects. Data are shown separately for each experimental condition (SiB versus SiSSN). Note that the unit for alpha and beta power is dB relative to one microvolt, calculated as 10log_10_(power/10^-12^).

Because alpha and beta power were both related to percent-correct score on a given trial, we directly compared alpha and beta power on a trial-by-trial basis. Figure 6A plots beta power versus alpha power in different trials across subjects and conditions, with data from each subject shown in a different color. We found significant correlation between alpha and beta power across trials [F(1,2081.5) = 175.7166, p < 2e-16], even after accounting for the random effect of subject (i.e., we see significant correlation even within subject). Another interesting observation from Figure 6A is that there appear to be individual differences in the overall magnitude of alpha [F(5,2093) = 1467.5081, p < 2e-16] and beta [F(5,2093) = 661.8108, p < 2e-16] power across trials, as well as in the distribution of the alpha-to-beta power ratio [F(5,2093) = 689.8876, p < 2e-16] across trials. The latter individual differences are quantified in Figure 6B. While some subjects (e.g., subject 46) show an alpha-to-beta power ratio greater than 1 across all trials, others (e.g., subject 64) show an alpha-to-beta power ratio less than 1 in most trials. This result raises the possibility that the listening strategy used may differ across individuals. However, our behavioral data do not allow us to directly address this possibility (see Discussion).

**Figure 6.**
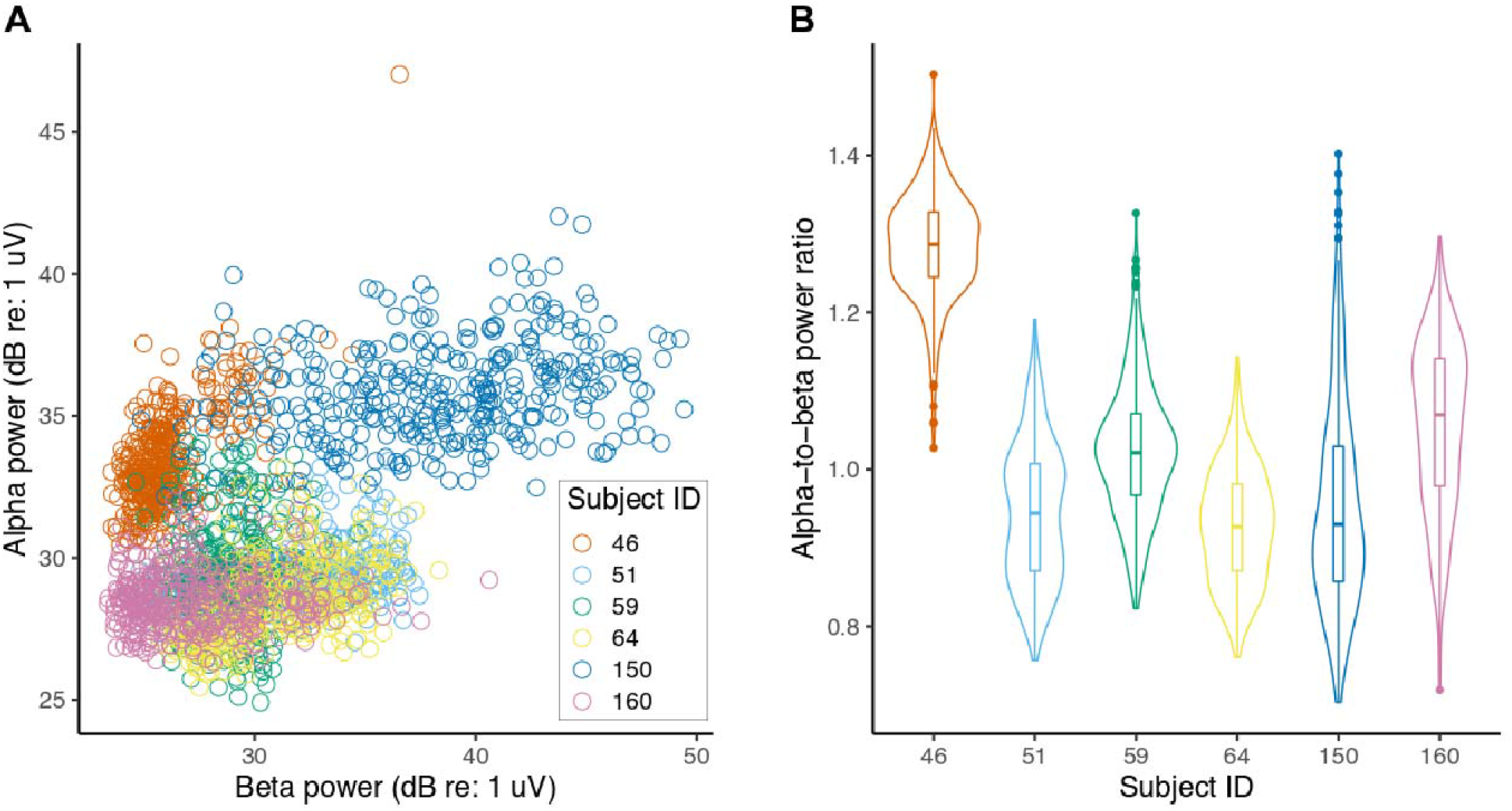
(A) Beta power versus alpha power in different trials (across subjects and conditions). The different colors in the plot correspond to different subjects. Note that the unit for alpha and beta power is dB relative to one microvolt, calculated as 10log_10_(power/10^-12^). (B) The distribution of alpha-to-beta power ratio across trials for each subject is shown as a violin plot. The median (horizontal bar), 50% confidence limits (box), and 95% confidence limits (whiskers) of each distribution are also shown.

Regardless of listening strategy, processes like selective attention and contextual prediction would be expected to affect the relationship between outcomes across different words in a trial in the average subject. To test whether this is the case, we plotted behavioral response error patterns. Figure 7 shows the histogram (across trials and subjects) of the number of keywords correct per sentence, separately for each experimental condition. These data show that the probability with which subjects get 0, 1, 2, 3, 4, or 5 keywords correct per sentence differs significantly (p-values shown in Figure 7; see Materials and Methods: Statistical analysis for details) from the expected distribution under the null hypothesis of independent keyword outcomes (i.e., if the performance on each keyword was not influenced by the others). Thus, there is a correlation in performance across the different keywords in a trial in both experimental conditions (SiB and SiSSN), which suggests a continuity effect consistent with top-down attention and contextual prediction. Note, however, that some bottom-up effects such as adaptation and deviance processing [56-58], and delayed top-down effects like postdiction [59, 60] may also contribute to the observed correlated outcomes across words.

**Figure 7.**
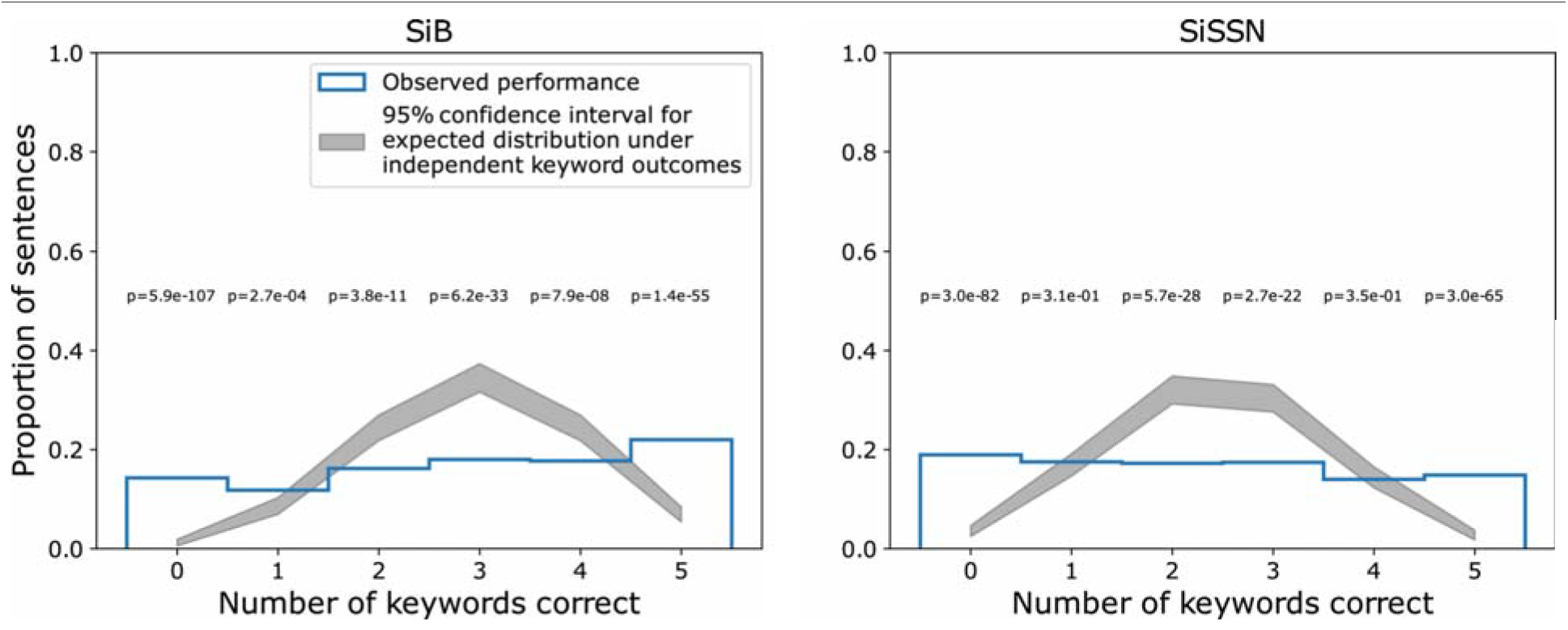
Normalized histogram (across trials and subjects) of number of keywords correct per sentence for SiB and SiSSN conditions (blue unfilled bars). Shown also is the 95% confidence interval for the expected distribution under the null hypothesis of independent keyword outcomes (gray patch). P-values for the observed data under the null hypothesis are also indicated.

Although alpha and beta power were correlated across trials (Figure 6A), beta power contributed significant additional predictive power to predict within-condition percent-correct score over the contribution of alpha power alone, and vice-versa (see Figure 8 and Table 1). Thus, not only do induced oscillations in both alpha and beta bands relate to speech intelligibility in noise on a trial-by-trial basis within condition, but crucially, alpha and beta power each make significant independent contributions to predicting trial outcome (i.e., trial-wise speech intelligibility).

**Figure 8.**
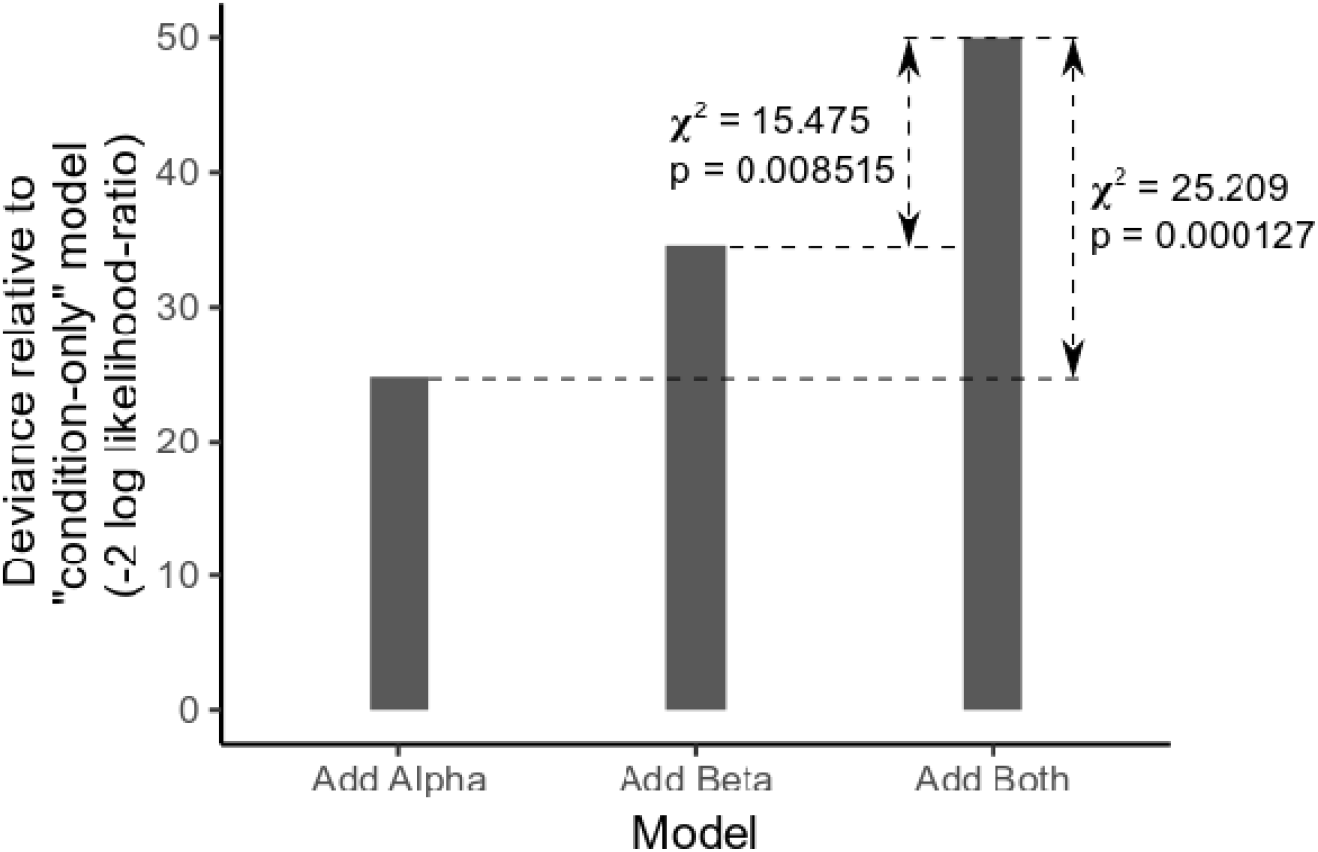
Likelihood ratio deviances (relative to a base model with condition as the only predictor) for three different multinomial models of percent-correct score: (1) model with condition and alpha power as predictors, (2) model with condition and beta power as predictors, and (3) model with condition, alpha power, and beta power as predictors. The deviance chi-square statistic and corresponding p-value are indicated for the comparison between models (1) and (3), and separately also for the comparison between models (2) and (3).

**Table 1.** Analysis of deviance table (Type II tests) for the multinomial linear regression analysis to test whether beta power contributes additionally to predicting percent-correct score over the contribution of alpha power alone, and vice-versa.

|  | Chi-square | Degree of freedom | Probability(>Chi-square) |
| --- | --- | --- | --- |
| alpha | 15.475 | 5 | 0.008515 |
| beta | 25.209 | 5 | 0.000127 |
| condition | 38.984 | 5 | 2.393e-07 |

Because of our observation that pre- and during-stimulus power were correlated for both alpha and beta, and that the average power across both periods covaries with single-trial behavioral outcomes, we wished to further understand the temporal evolution of the two rhythms over the time course of the trial. While it is well-established that alpha enhancement begins in the preparatory period (for example, before stimulus onset but after cueing subjects to “stay still listen now” as in our study), the temporal dynamics of the beta rhythm during speech perception in noise is not as well studied. Thus, we contrasted the scalp topographic maps between the during- and pre-stimulus periods for both beta and alpha (Figure 9) to obtain further insight. Figure 9 shows that parieto-occipital alpha power is stronger in the during-stimulus period (Panel A), consistent with maintaining an increasing attentional focus on the target speech. However, the scalp topomap difference between during- and pre-stimulus periods in the beta band (Panel B) shows regions of reduced power fronto-centro-laterally in both hemispheres, and regions of increased power fronto-medially. This suggests that the beta power observed in the present study consists of two functionally distinct components. The fronto-centro-lateral component of beta, which is stronger in the pre-stimulus period (i.e., before stimulus onset but after the “stay still and listen now” cue), is consistent with motor suppression during attentive engagement [61, 62]. The fronto-medial component that is stronger during stimulus presentation may be associated with predictive coding mechanisms [22–24, 30]. Given this temporal evolution of alpha and beta, we examined the association between pre-stimulus power and performance and separately between during-stimulus power and performance. For both alpha and beta, pre- and during-stimulus power were each significantly associated with single-trial percent-correct score within condition [Supplementary Figures S1 and S2; F(5,2093) = 4.1434, p = 0.0009513 for pre-stimulus alpha; F(5,2093) = 4.7789, p = 0.0002397 for during-stimulus alpha; F(5,2093) = 5.6198, p = 3.753e-05 for pre-stimulus beta; F(5,2093) = 4.7979, p = 0.00023 for during-stimulus beta), as might be expected given the high correlation between pre- and during-stimulus power (Figure 4).

**Figure 9.**
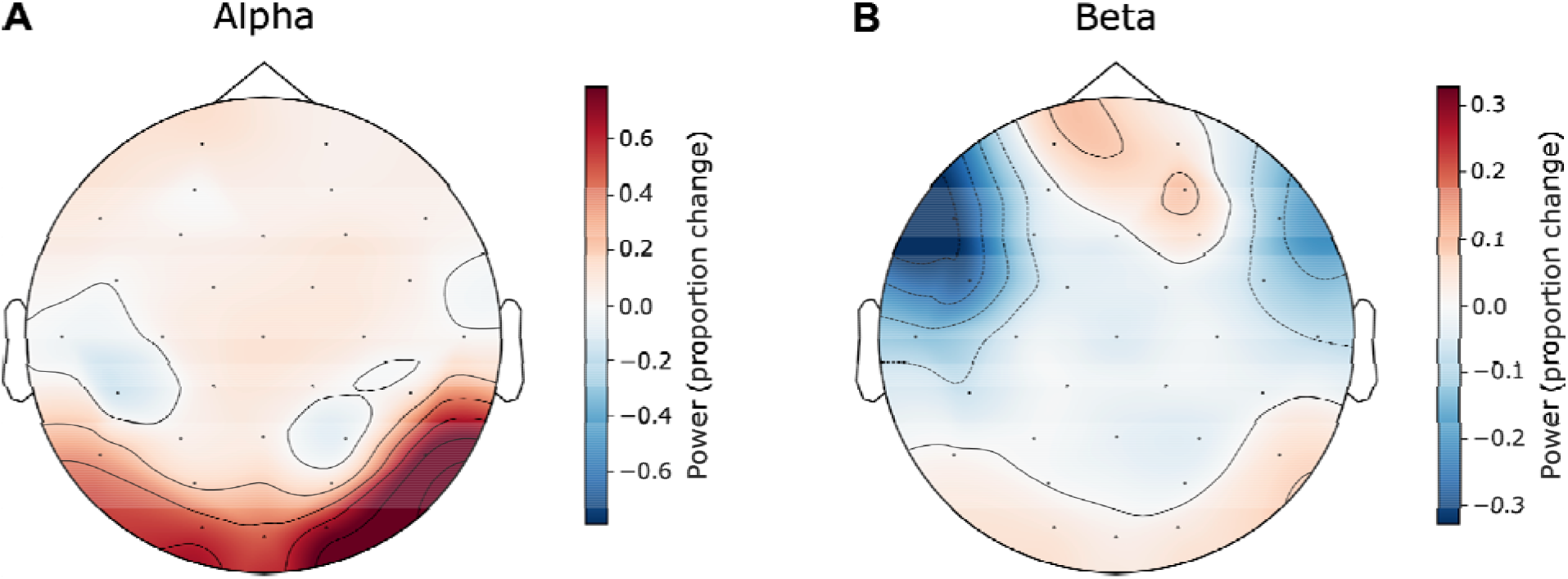
Proportion change in the average scalp topography map for alpha (Panel A) and beta (Panel B) from the pre- to the during-stimulus period [(during-stimulus - pre-stimulus)/pre-stimulus].

## 5. Discussion

Using human EEG with simultaneous speech intelligibility measurements in different masking conditions (speech in multi-talker babble, and speech in speech-shaped noise) in the present study, we found that induced brain oscillations in the alpha and beta bands relate to speech intelligibility in competition on a trial-by-trial basis. Specifically, we found that the overall magnitudes (averaged over the pre- and during-stimulus periods) of alpha power in parieto-occipital EEG channels and beta power in frontal channels significantly covary with, and importantly independently contribute to, single-trial speech intelligibility in our speech-in-noise tasks (Figures 5, 8). These results are consistent with the posited role of the parieto-occipital alpha rhythm in auditory selective attention [10–14, 16–18] and the frontal beta rhythm in maintenance of the current sensorimotor state [22] and sensorimotor predictive coding [23, 24] that is thought to stabilize speech representation in adverse listening conditions [25–29]. The interpretation that some combination of these top-down effects influences single-trial behavioral outcome is also supported by the observed correlation in performance across words within a trial (Figure 7).

Our results are in line with prior reports of a positive correlation between alpha power in parietal EEG channels and speech intelligibility in noise (e.g., across SNRs as quantified in the during-stimulus period by Hall et al. [20], and across individuals as quantified in the pre-stimulus period by Alhanbali et al. [21]). However, at least at first glance, our results appear to be at odds with other reports (e.g., by Obleser and Weisz [63] and Becker et al. [64], who used noise-vocoded speech in quiet, and Dimitrijevic et al. [65], who used digits in noise) that better comprehension is associated with alpha suppression (rather than a power increase) in the late during-stimulus period in temporal brain regions and central EEG channels. This discrepancy may be explained in part by the existence of multiple neural generators of task-related alpha (i.e., alpha power in the parieto-occipital and central EEG channels may reflect two different mechanisms of alpha [18]). Moreover, some of these studies presented speech in quiet rather than with simultaneous competing sounds, which could evoke different mechanisms [63, 64].

Foxe and Snyder [10] distinguish between parieto-occipital alpha seen in an unaroused state (e.g., when visual stimuli are ignored [15]) and that seen in selective attention across different stimuli (especially spatial selective attention, where alpha power is lateralized according to the hemifield of focus; [16–18]). In the present study, the target speech and masker sources were both presented diotically rather than spatially separated; thus, even though it required selective attention, our task did not involve any spatial focus of attention. It may be that the alpha in the current study, which covaries with trial-wise speech intelligibility, reflects an overall suppression of the visual scene and focus of auditory attention, rather than a mechanism specific to stimulus selection. Another possibility is that there may be a common mechanism in play across the parieto-occipital alpha seen in the two cases. Indeed, the frontoparietal attention network becomes active during spatial attention and working memory for auditory stimuli as well as for visual inputs, even though many earlier studies assume it is strictly a visuospatial processing network [66–70]. Thus, future studies should disambiguate between the different mechanisms by which the alpha rhythm may mediate suppression of sensory distractors [10–14], especially for co-localized sources like those used in the current study.

Unlike parieto-occipital alpha, the functional role of frontal beta in auditory perception is less understood. That parieto-occipital alpha is associated with attentional focus and is present even before stimulus onset (Figures 2, 3A) is well documented [10–19]. However, in the current study we find that frontal beta power in both the pre-stimulus and during-stimulus periods covaries with single-trial speech-in-noise outcomes (Supplementary Figures S1 and S2; statistics given in Results). Although our results about during-stimulus beta may potentially be explained by invoking the predictive coding theory [22–24, 30], the role of pre-stimulus frontal beta is less clear. The scalp topomap result shown in Figure 9B suggests that the beta power observed in the present study consists of two functionally distinct components. The fronto-centro-lateral component is stronger in the pre-stimulus period and may reflect a mechanism that suppresses neuronal processing of new movements, favoring maintenance of the current sensorimotor state [22, 61, 62, 71–74]; in the present study this motor suppression may begin as subjects prepare for the upcoming stimulus after being cued to “stay still and listen now”. In contrast, the fronto-medial component of beta may be a network mechanism spanning fronto-motor and auditory areas for top-down prediction/anticipation that may be active during both pre- and during-stimulus periods [22–24, 30, 75]. Our behavioral manipulations and 32-channel EEG recordings cannot further disambiguate between these two components of beta. Nevertheless, we find that although pre-stimulus beta covaried with during-stimulus beta and during-stimulus power levels were similar to pre-stimulus levels (Figure 4B), during-stimulus frontal beta power contributed significant additional predictive power to predict within-condition percent-correct score over the contribution of pre-stimulus power alone (Supplementary Table S2). Future experiments should be designed to dissociate beta rhythms associated with motor suppression during attentional engagement from beta activity associated with dynamic predictive coding mechanisms. In particular, high-density EEG or magnetoencephalography (MEG) recordings along with source-space analysis can be used to probe which specific beta-band mechanisms relate to speech understanding in competition (e.g., beta-band synchrony between auditory and fronto-motor areas would imply a different mechanism from beta activity that is confined to motor cortex).

Our current results show that trial-by-trial variations in alpha and beta power are correlated, even within subject (Figure 6A; statistics given in Results). Prior studies have also reported that oscillatory activity within the alpha and beta bands are correlated [76], even though they may represent distinct functions. Despite being correlated, alpha and beta power each provide significant independent contributions to predicting single-trial percent-correct score (Figure 8). Moreover, there are individual differences in the overall magnitude of alpha and beta power across trials (Figure 6A). Comparing these neural individual differences (Figure 6A) to the individual differences in behavioral performance (Figure 1) leads us to hypothesize that a greater alpha or beta power for an individual subject might relate to greater average performance for that subject; however, we are unable to test this specific hypothesis due to the low statistical power (just 6 subjects) in our study to conduct such an analysis of individual differences. Rather, we relate trial-by-trial fluctuations in alpha and beta power to trial-wise variations in behavioral outcome. Our data (Figure 6B) suggests that there may also be individual differences in the alpha-to-beta power ratio across trials, which raises the possibility that listeners used different task strategies. Regardless of listening strategy used, trial-wise performance improved as either alpha or beta power increased (Figure 5). Our behavioral measurements cannot elucidate the specific listening strategy used by any particular subject (e.g., focused attention or predictive coding). However, measuring confusion patterns can inform which listening strategy an individual subject used (e.g., when a subject made an error, whether they reported a word from the competing stream or a new, contextually suitable word [77]). This approach should be explored in future experiments.

## Acknowledgments

This study was supported by grants from the National Institutes of Health [Grant Nos. F31DC017381 (to V.V.), T32DC11499 (to V.V.), F32DC020649 (to V.V.), R01DC009838 (to M.G.H.), R01DC015989 (to H.M.B.), R01DC013825 (to B.G.S.-C.), and R01DC019126 (to B.G.S.-C.)] and from Action on Hearing Loss [Grant No. G72 (to M.G.H.)].

