## Supplementary Information for "Induced Alpha And Beta Electroencephalographic Rhythms Covary With Single-Trial Speech Intelligibility In Competition"

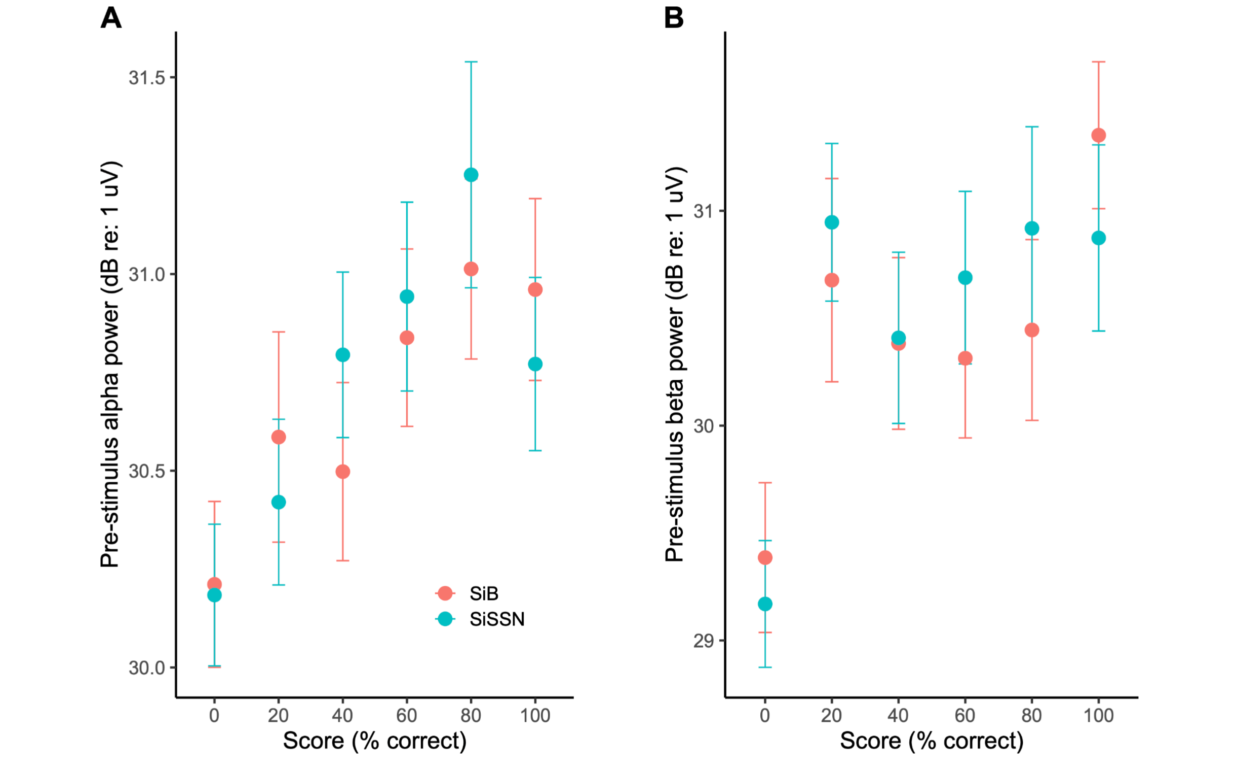


**Supplementary Figure S1.** Pre-stimulus alpha (A) and pre-stimulus beta (B) power [mean and standard error of the mean (STE)] versus percent-correct score in different trials across subjects. Data are shown separately for each experimental condition (SiB versus SiSSN).

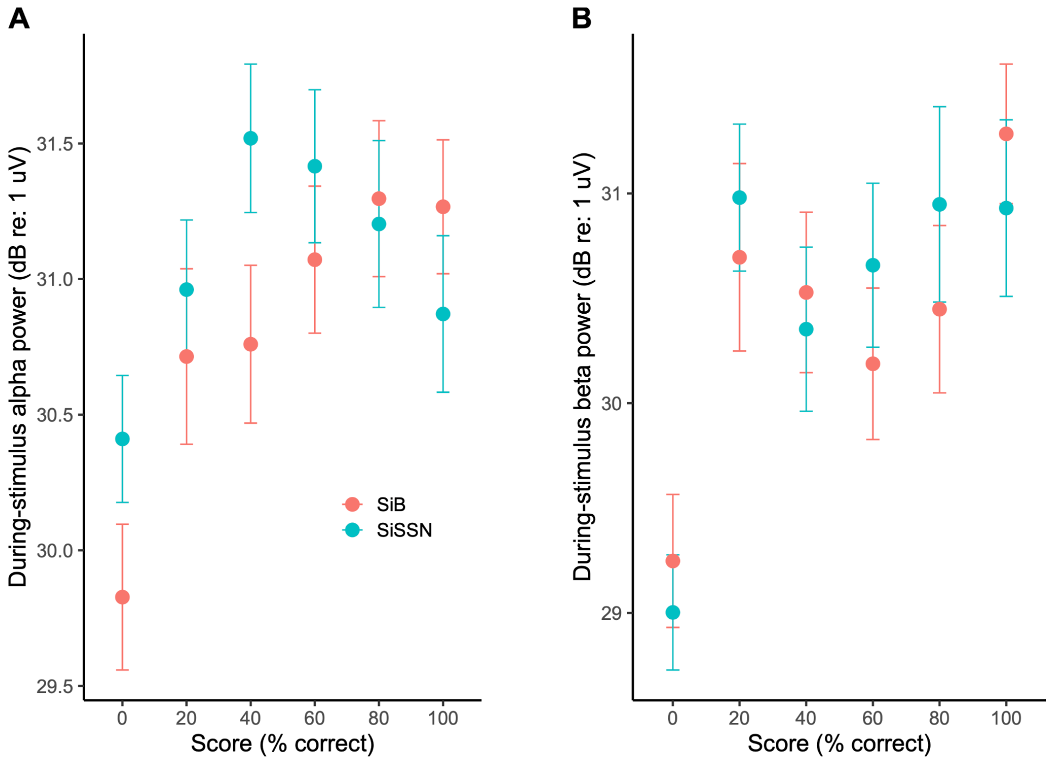


**Supplementary Figure S2.** During-stimulus alpha (A) and during-stimulus beta (B) power (mean and STE) versus percent-correct score in different trials across subjects. Data are shown separately for each experimental condition (SiB versus SiSSN).

**Supplementary Table S1.** Analysis of deviance table (Type II tests) for the multinomial linear regression analysis to test whether during-stimulus alpha power contributes additionally to predicting percent-correct score over the contribution of pre-stimulus alpha power alone, and vice-versa.

R code (uses the “nnet” package):

```
model <- multinom(formula = percentcorrect ~ duringstimulus_alpha + prestimulus_alpha + condition,
data = data)
Anova(model)
```

| | Chi-square | Degree of freedom | Probability(>Chi-square) |
|---|---|---|---|
| duringstimulus_alpha | 13.912 | 5 | 0.01618 |
| prestimulus_alpha | 9.933 | 5 | 0.07717 |
| condition | 38.673 | 5 | 2.763e-07 |

**Supplementary Table S2.** Analysis of deviance table (Type II tests) for the multinomial linear regression analysis to test whether during-stimulus beta power contributes additionally to predicting percent-correct score over the contribution of pre-stimulus beta power alone, and vice-versa.

R code (uses the “nnet” package):

```
model <- multinom(formula = percentcorrect ~ duringstimulus_beta + prestimulus_beta + condition,
data = data)
Anova(model)
```

| | Chi-square | Degree of freedom | Probability(>Chi-square) |
|---|---|---|---|
| duringstimulus_beta | 12.931 | 5 | 0.02404 |
| prestimulus_beta | 4.089 | 5 | 0.53674 |
| condition | 38.371 | 5 | 3.178e-07 |
